# Dysregulation of NRAP degradation by KLHL41 contributes to pathophysiology in Nemaline Myopathy

**DOI:** 10.1101/487454

**Authors:** Caroline Jirka, Jasmine H Pak, Claire A Grosgogeat, Michael Mario Marchetii, Vandana A Gupta

**Affiliations:** Division of Genetics, Brigham and Women’s Hospital, Harvard Medical School, Boston, MA-02115

**Keywords:** Nemaline Myopathy, Skeletal muscle, Ubiquitination, E3 Ligase, Cul3, Thin filament Chaperone, NRAP, zebrafish, Myofibrillogenesis, Downregulation, Therapy

## Abstract

Nemaline myopathy (NM) is the most common form of congenital myopathy that results in hypotonia and muscle weakness. This disease is clinically and genetically heterogeneous, but three recently discovered genes in NM encode for members of the Kelch family of proteins. Kelch proteins act as substrate-specific-adapters for CUL3 E3 ubiquitin ligase to regulate protein turn-over through the ubiquitin-proteasome machinery. Defects in thin filament formation and/or stability are key molecular processes that underlie the disease pathology in NM, however, the role of Kelch proteins in these processes in normal and diseases conditions remains elusive *in vivo*. Here, we describe a role of NM causing Kelch protein, KLHL41, in premyofibil-myofibil transition during skeletal muscle development through a regulation of the thin filament chaperone, NRAP. KLHL41 binds to the thin filament chaperone NRAP and promotes ubiquitination and subsequent degradation of NRAP, a process that is critical for the formation of mature myofibrils. KLHL41 deficiency results in abnormal accumulation of NRAP in muscle cells. NRAP overexpression in transgenic zebrafish resulted in a severe myopathic phenotype and absence of mature myofibrils demonstrating a role in disease pathology. Reducing Nrap levels in KLHL41 deficient zebrafish rescues the structural and function defects associated with disease pathology. We conclude that defects in KLHL41-mediated ubiquitination of sarcomeric protein contribute to structural and functional deficits in skeletal muscle. These findings further our understanding of how the sarcomere assembly is regulated by disease causing factors *in vivo*, which will be imperative for developing mechanism-based specific therapeutic interventions.

## Introduction

Muscle development is a multistep process that involves the specification of myogenic progenitors and myoblasts and subsequent fusion of myoblasts that generates syncytial, multinucleated myotubes (1-3). Muscle cells typically contain dozens of myofibrils, each consisting of many sarcomeres, the smallest functional contractile unit of muscle. Consequently, mutations in genes encoding proteins that are critical for sarcomere assembly, and efficient coupling to the excitation-contraction machinery are associated with several forms of myopathies in humans (4-9). Despite a reasonable understanding of the mature sarcomere and myofibrils architecture, the assembly mechanism of protein complexes that leads to myofibril and sarcomere formation during muscle development remains less understood.

The process of sarcomeric assembly is mostly studied during myoblasts differentiation or in cardiomyocytes *in vitro* and has led to several models of sarcomere assembly with a common premise that shows premyofibrils formation begins with the polymerization of actin monomers into thin filaments by Arp2/3 protein complex whose barbed ends (fast-growing) are aligned and crossed linked by a-actinins in Z-lines, the borders of sarcomeres (3, 10, 11). The pointed ends (slow-growing) extend towards the M-line, where they interdigitate with thick filaments and form premyofibrils. Mature myofibrils are formed through lateral fusion of premyofibrils with addition of proteins that stabilize the core and structure of developing myofibrils (12). While the role of a few central structural proteins during sarcomere assembly is established, factors and mechanism that regulate the dynamics and transition between different stages of myofibrillogenesis remains elusive. Recent studies have identified mutations in a number of genes encoding non-structural proteins contributing to defects in sarcomeric architecture and muscle weakness in skeletal muscle disorders (13-18). Determining the mechanism of sarcomeric assembly *in vivo* during vertebrate myogenesis by these factors is crucial for identifying the critical processes that control the muscle development, and ultimately for determining how disease-causing variants in these genes contribute to muscle weakness in human myopathies.

Nemaline myopathy (NM) is a rare congenital disorder primarily affecting skeletal muscle function (19-21). Clinically, NM is a heterogeneous group of myopathies of variable severity. The “severe” congenital form of NM presents with reduced or absent spontaneous movements *in utero* leading to severe contractures or fractures at birth and respiratory insufficiency leading to early mortality. NM is a genetically heterogeneous condition, and mutations in 11 different genes have been identified by us and others that are associated with dominant and/or recessive forms of this disease (4, 5, 15, 22-29). While eight genes encode structural components of thin filaments in sarcomeres, the function of newly discovered genes encoding Kelch proteins-KBTBD13, KLHL40 and KLHL41 in muscle development and function remains to be elucidated. Recent *in vitro* studies have shown that these Kelch proteins act as substrate specific adapters for Cul3-E3 ubiquitin ligases that regulate the stability and turnover of specific target proteins by ubiquitination (30-33).

The protein turn-over process is highly dynamic and provides a mechanism for the removal of unwanted proteins or changes in protein isoforms without disassembling the structural integrity needed for the myofibril function (34). The proteome fidelity is mediated by the ubiquitin-proteasome system (UPS), which constitutes the central protein-turnover machinery in cells. The specificity to UPS in development, diseases and, more importantly, tissue-specific context is provided by E3 ubiquitin ligases. E3 ubiquitin ligases initially catalyze ubiquitin transfer from an E2-ligase to their target substrates and subsequent polyubiquitination from various linkage-specific E2s (35). One subgroup of E3 ligases are the ubiquitously expressed cullins that do not bind to their substrates directly but rely on an array of adapter proteins such as Kelch proteins (36, 37). Ubiquitination-mediated protein turnover has been the focus of intense investigation in the catabolic processes contributing to muscular atrophies and degeneration (38-41). On the contrary, there is very limited knowledge available about the ubiquitin-dependent machinery during skeletal muscle development and how such machinery regulates different aspects of myogenesis. As mutations in several Kelch protein encoding genes result in skeletal muscle diseases, an understanding of the role of Kelch proteins in ubiquitination-mediated protein turn-over during skeletal muscle development may provide novel therapeutic intervention in NM and related diseases.

Our current work has addressed these crucial points to understand the mechanism of KLHL41-mediated ubiquitination of a thin filament chaperone, NRAP, on regulating the skeletal muscle structure, function and disease pathogenesis of nemaline myopathy. We show that KLHL41 interacts with and destabilizes NRAP, a protein that is critical for premyofibril-myofibril transition. KLHL41 deficiency results in the accumulation of NRAP protein which leads to myopathy in zebrafish. In KLHL41 deficiency, overexpression of NRAP prevents binding of KLHL40 to nebulin that is required to stabilize thin filaments in mature myofibers. Finally, downregulation of NRAP in KLHL41 deficiency improves disease pathophysiology. These studies highlight the role of KLHL41-regulated ubiquitination in skeletal muscle development and provide new insights on disease pathology in Nemaline Myopathy.

## Results

### Identification of *in vivo* substrates for CUL3-adapter protein, KLHL41 in skeletal muscle

Cullin 3 ubiquitin ligase (CUL3) requires BTB-Kelch proteins as substrate specific adapters to direct the ubiquitination of target proteins. A previous *in vitro* study has demonstrated that KLHL41-CUL3 interaction is crucial for CUL3-mediated protein ubiquitination of substrate proteins (33). Kelch proteins interact with their substrates for ubiquitination through Kelch domains (47). Therefore, to identify the molecular interactors of KLHL41 in skeletal muscle and gain insights into pathophysiological mechanism of NM, we performed a yeast two-hybrid screening (Y2H) using the Kelch domain of KLHL41 protein as a bait against a human fetal and adult skeletal muscle library and identified 350 clones coding nebulin (NEB), nebulin related anchoring protein (NRAP), Proteasome Maturation Protein (POMP), interferon-related developmental regulator 1 (IRFD1) and RNA Binding Motif Protein 4 (RBM4) (Fig. 1A and Table S1). Nemaline myopathy is primarily a thin filament disease (48). Recent work has shown that KLHL41 interacts with nebulin to stabilize thin filaments (33). However, implications of KLHL41-NRAP interactions *in vivo* are not known. Therefore, we focused our studies on the role of KLHL41-NRAP in skeletal muscle development and maintenance using nebulin as a positive control. Sequencing of different interacting clones reveled that KLHL41 interacted with nebulin super repeats in NRAP protein. KLHL41 also interacted with nebulin through interactions with different domains: nebulin repeats, nebulin super repeats, C-terminal serine rich region and SH3 domain (Fig. 1A).

**Figure 1.**
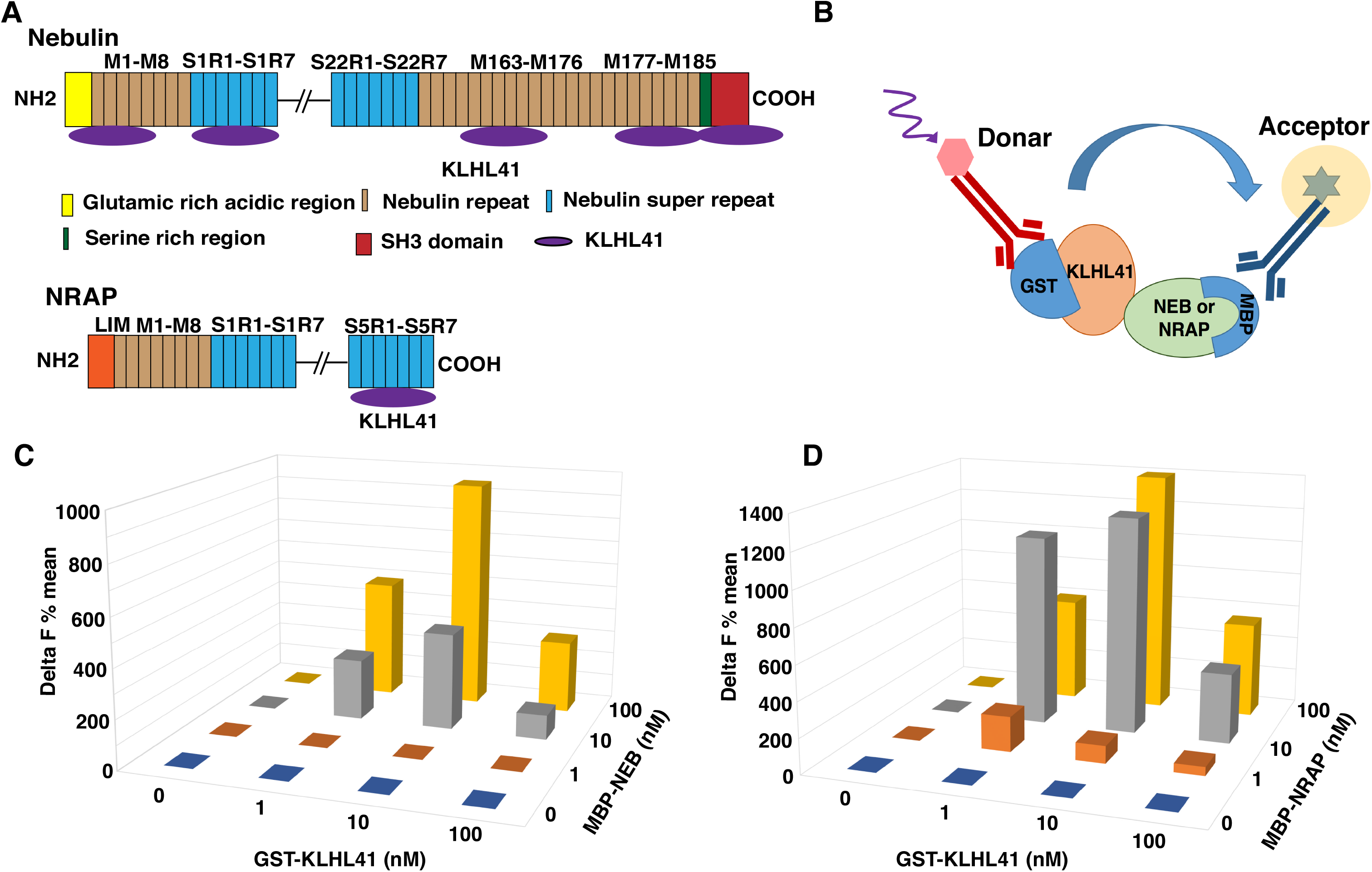
KLHL41 protein interactors in skeletal muscle. (A) Schematics of regions on nebulin and NRAP proteins that interact with KLHL41, identified by yeast 2 hybrid screening (B) Principle of the HTRF method for detecting interaction between KLHL41-GST and Nebulin or NRAP-MBP fusion proteins. Antibodies against GST or MBP tags were labeled donor or acceptor, respectively. The HTRF signals are generated on close proximity of each of the labeled antibodies through direct interaction of two proteins (C) Quantification of the interaction between different concentrations of KLHL41 and nebulin and (D) NRAP interactor proteins; where KLHL41 is GST tagged and nebulin or NRAP are MBP tagged.

To validate these interactions, we employed homogeneous time-resolved fluorescence (HTRF). The HTRF technology is based on time resolved fluorescence energy transfer (TR-FRET) that occurs between long-lived fluorophore Europium or Terbium as a donor and d2 as acceptors (Fig. 1B). The HTRF interaction between two proteins is quantified by calculating delta F (%) (described in methods) (49). Different interacting fragments of NEB and NRAP were expressed as MBP fusion proteins in BL21 *E. coli* cells and purified using affinity purification. Three different concentrations of interacting proteins (1, 10 and 100 nM) were tested in the HTRF assay that confirmed the interactions of KLHL41 with NEB and NRAP (Fig.1C-D). Under these conditions, best delta F% signal detected for KLHL41-NEB was 950 (10nM KLHL41 and 100nM NEB). The highest delta F% signal detected for KLHL41 (10nM) and NRAP (100mM) was 1390 (10nM KLHL41 and 100nM NRAP). Higher delta F% values were observed for KLHL41-NRAP interaction compared to KLHL41-NEB interactions suggesting a higher affinity of KLHL41-NRAP interactions in comparison to KLHL41-NEB interactions. As protein-protein interactions identified by yeast two hybrid assay are independent of post-translational modifications, our studies demonstrate that KLHL41 interacts with nebulin and NRAP through direct protein-protein interactions.

### KLHL41 regulates the stability of thin-filament chaperone NRAP by ubiquitination

To validate the interaction of KLHL41 and NRAP in myoblasts, KLHL41-FLAG and NRAP-V5 were over-expressed in C2C12 myoblasts, and cell extracts were immunoprecipitated by an antibody against V5 (Fig. 2A). This resulted in co-immunoprecipitation of KLHL41-FLAG with NRAP-V5 but not with control mouse IgG, confirming that KLHL41 interacts with NRAP in the myogenic context. Kelch proteins interact with protein substrates for ubiquitination through E3 Cul3 ligases (47). Therefore, we hypothesized these protein-protein interactions may be required to regulate ubiquitination of the interacting partners such as NRAP. To investigate the ubiquitnation of NRAP by KLHL41, we overexpressed human KLHL41-FLAG with NRAP-V5 plasmids in C2C12 myoblasts in the presence of ubiquitin-HA plasmid. The amounts of KLHL41 plasmid was varied (0-1.0 ng) whereas the amount of NRAP protein was kept constant (2.0 ng) (Fig. 2B). 48 hours post transfections, NRAP was immuno-precipitated with a V5 antibody and analyzed by western blot analysis. Western-blot analysis demonstrated that ubiquitination of NRAP was increased with an increase in KLHL41 concentration. Interestingly, at higher concentrations of KLHL41, decreased levels of ubiquitinated NRAP protein was observed. These results suggest that KLHL41-mediated ubiquitination resulted in the degradation of NRAP, potentially through proteolysis by the proteasome degradation pathway.

**Figure 2.**
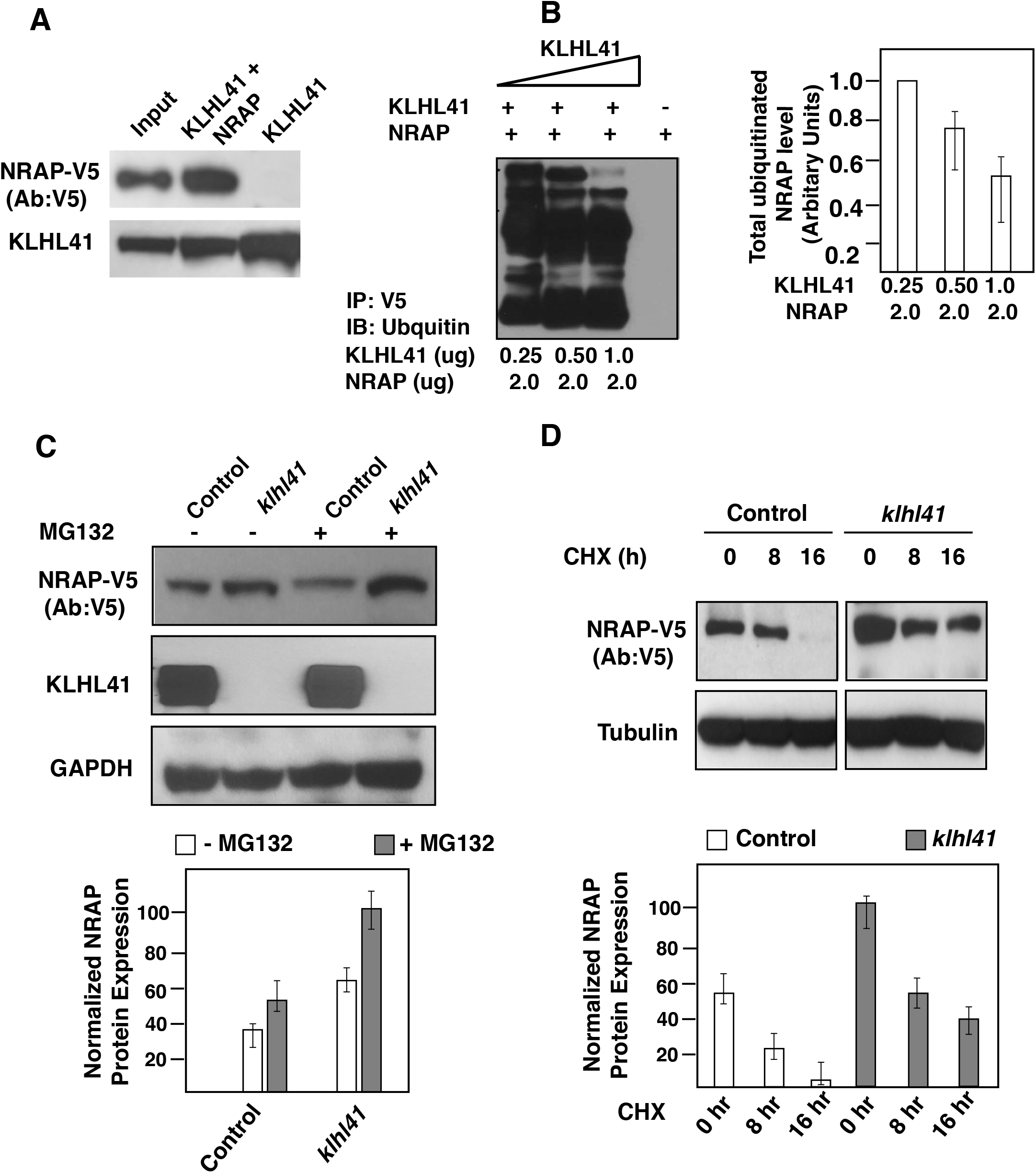
KLHL41-NRAP interaction results in ubiquitination and proteasome-mediated degradation of NRAP. (A) Co-immunoprecipitation in C2C12 cells transfected with KLHL41-FLAG and NRAP-V5 demonstrating interaction of the two proteins in the myogenic context (B) Overexpression of KLHL41-FLAG and NRAP-V5 in the presence of ubiquitin-HA in C2C12 cells results in increased ubiquitination and decreased stability of NRAP (C) Western blot analysis and quantification of NRAP in control or *Klhl41* knockout myoblasts treated with MG132 (D) Western blot analysis and quantification of NRAP in control or *Klhl41* knockout myoblasts treated with cycloheximide for different time intervals.

### KLHL41 deficiency result in increased NRAP protein

As KLHL41 is required for the ubiquitination and subsequent degradation of NRAP, we investigated the effect of KLHL41 deficiency on NRAP levels. To study the effect of KLHL41 deficiency on NRAP in a mammalian disease model, we created *Klhl41* mouse C2C12 knockout cell lines using CRISPR technology. *Klhl41* knockout myoblasts exhibited increased levels of NRAP protein in comparison to control cells as observed by Western blot (Fig. 2C). No changes in *Nrap* gene transcription were observed by q-PCR, validating that the effect of KLHL41-mediated degradation of NRAP is regulated at the post-translational level. Consistent with the role of ubiquitin-mediated proteasomal degradation, NRAP downregulation was mitigated by proteasome inhibitor, MG132, in control cells (Fig. 2C). The role of endogenous KLHL41 on endogenous NRAP protein turnover was further investigated by cycloheximide chase assay in control and *Klhl41* knockout cells (Fig. 2D). Control cells exhibited increased turnover of NRAP, and no protein was detected after 24 hours of cycloheximide treatment. Conversely, *Klhl41* knockout cells exhibited a reduced turnover of NRAP protein in comparison to control cells, and a significant amount of NRAP protein accumulation was still observed after 24 hours of the cycloheximide chase. Collectively, these data indicate that KLHL41 controls NRAP protein levels in skeletal muscle by ubiquitination-mediated proteasomal degradation.

### NRAP accumulation leads to myopathy in zebrafish

As KLHL41 deficiency results in accumulation of NRAP in C2C12 *Klhl41* myoblasts, we investigated the *in vivo* impact of NRAP accumulation on skeletal muscle structure and function. We created a transgenic *NRAP* overexpressing zebrafish under the control of skeletal muscle specific myosin promoter (*mylz*) (Fig. 3A)(50). Human *NRAP* was integrated in to zebrafish genome utilizing the Tol2 transposon-based system (46). Genomic integration and transgene expression were monitored using a GFP reporter-protein linked to the coding sequence of *NRAP* (*mylz:NRAP-gfp*). Germline transmission of the *NRAP* transgene was confirmed by PCR-based genotyping. Additionally, the expression of hu-NRAP-GFP protein was confirmed by immunoblotting with a GFP antibody at 3 dpf that showed ~2.4 folds upregulation of the transgene compared to wild-type controls (Fig. 3J). To investigate if NRAP overexpression affects the structure and function of skeletal muscle, transgenic zebrafish larvae were analyzed at 3 dpf by brightfield microscopy. Zebrafish larvae expressing NRAP transgene demonstrated extensive hypotonia and dorsal curvature indicative of a myopathic phenotype (Fig. 3B-C). To test if the morphological abnormalities in the *mylz:NRAP-gfp* transgenic fish are a consequence of structural defect in skeletal muscle, birefringence analysis was performed. Visualization of *mylz:NRAP-gfp* transgenic fish under polarized lenses showed a significant reduction in birefringence in comparison to control indicative of structural disorganization of skeletal muscle (Fig 3. D-E). To determine the specific defects in skeletal muscle structure due to NRAP overexpression, immunofluorescence and electron microscopy were performed in the *mylz:NRAP-gfp* transgenic and control zebrafish (3dpf). Whole mount immunofluorescence of skeletal muscle by phalloidin staining revealed thinner myofibers and gaps between adjacent myofibers in *mylz:NRAP-gfp* zebrafish (Fig. 3F-G). Additionally, visualization of skeletal muscle ultrastructure by electron microscopy revealed smaller and fewer sarcomeres in the *mylz:NRAP-gfp* transgenic fish in comparison to the control. While assembly of immature actin-myosin bundles was observed, a lack of mature myofibrils was evident in the *mylz:NRAP-gfp* transgenic zebrafish in comparison to the control (Fig. 3H-I, arrow). Evaluation of the skeletal muscle function by swimming behavior analysis also revealed a blunted motor activity in *mylz:NRAP-gfp* transgenic fish compared to controls (Fig. 3K). Together, these data provide evidence that overexpression of NRAP prevents the formation of mature myofibers and results in reduced motor function in affected skeletal muscle.

**Figure 3.**
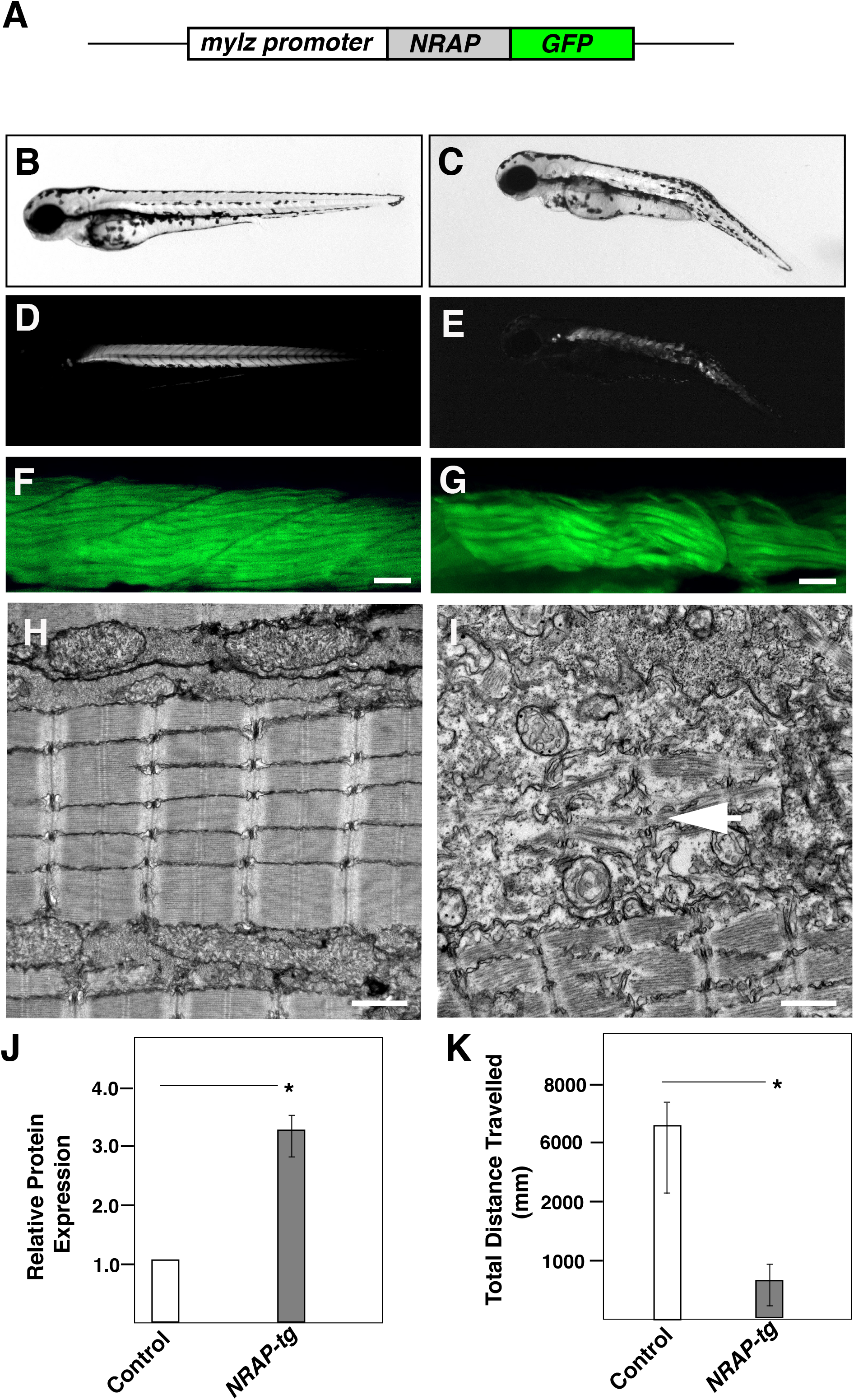
Transgenic NRAP zebrafish exhibit structural and functional deficits. (A) Schematics of NRAP transgenic construct. Human NRAP cDNA was cloned inframe with GFP reporter and downstream of the zebrafish myosin light chain promoter using tol2-transgenesis (B-C) *Tg* (*mylz:NRAP-gfp*) fish exhibit dorsal curvature and leaner bodies indicative of a myopathic phenotype (3dpf) (D-E) Visualization of *Tg* (*mylz:NRAP-gfp*) under polarize light revealed reduced birefringence compared to control (3 dpf) (F-G) Whole mount phalloidin staining of *Tg* (*mylz:NRAP-gfp*) larvae (3 dpf) exhibiting fewer myofibers and smaller myotomes in comparison to controls. Scale bar: 50μm (H-I) Transmission electron microscopy demonstrating mature myofibers in control fish whereas *Tg* (*mylz:NRAP-gfp*) fish exhibit presence of immature sarcomeres (white arrow) and lacked matured myofibrils (arrow). Scale bar: 500 nm (J) Quantification of the NRAP protein in transgenic zebrafish by western blot analysis with Gfp antibody (K) Quantification of the distance traveled by control and *Tg* (*mylz:NRAP-gfp*), acquired by an automated infra-red imager.

### NRAP prevents KLHL40-Nebulin interaction in KLHL41 deficiency

During skeletal muscle development, NRAP depletion corresponds to lateral fusion of premyofibrils and assembly of proteins complexes by scaffolding on KLHL41 resulting in mature thin filaments in sarcomeres. In addition to KLHL41, closely related NM causing Kelch protein, KLHL40, interacts with nebulin and NRAP in skeletal muscle and stabilizes mature thin filaments (32). Therefore, in KLHL41 deficiency, we investigated the impact of NRAP on complex formation between KLHL40 and nebulin to evaluate the functional redundancy between closely related KLHL40 and KLHL41. Co-expression of equimolar amounts of KLHL40-FLAG, NEB (C-terminal or N-terminal) and NRAP was performed in *klhl41* deficient C2C12 myoblasts (Fig. 4A-B). Immunoprecipitation of KLHL40-FLAG with FLAG antibody and western blotting revealed that KLHL40 co-immunoprecipitated with either NRAP or NEB, validating the interaction between these proteins. Interestingly, in the presence of both NRAP and NEB, KLHL40 preferentially co-immunoprecipitated with NRAP, and no interaction of KLHL40 was observed with NEB. These studies suggest that NRAP exhibit a high affinity for proteins, such as KLHL40 and KLHL41, that are essential for the assembly of proteins required in the formation of mature myofibrils, and the removal of NRAP by KLHL41, therefore, is a critical step in the formation of mature skeletal muscle.

**Figure 4:**
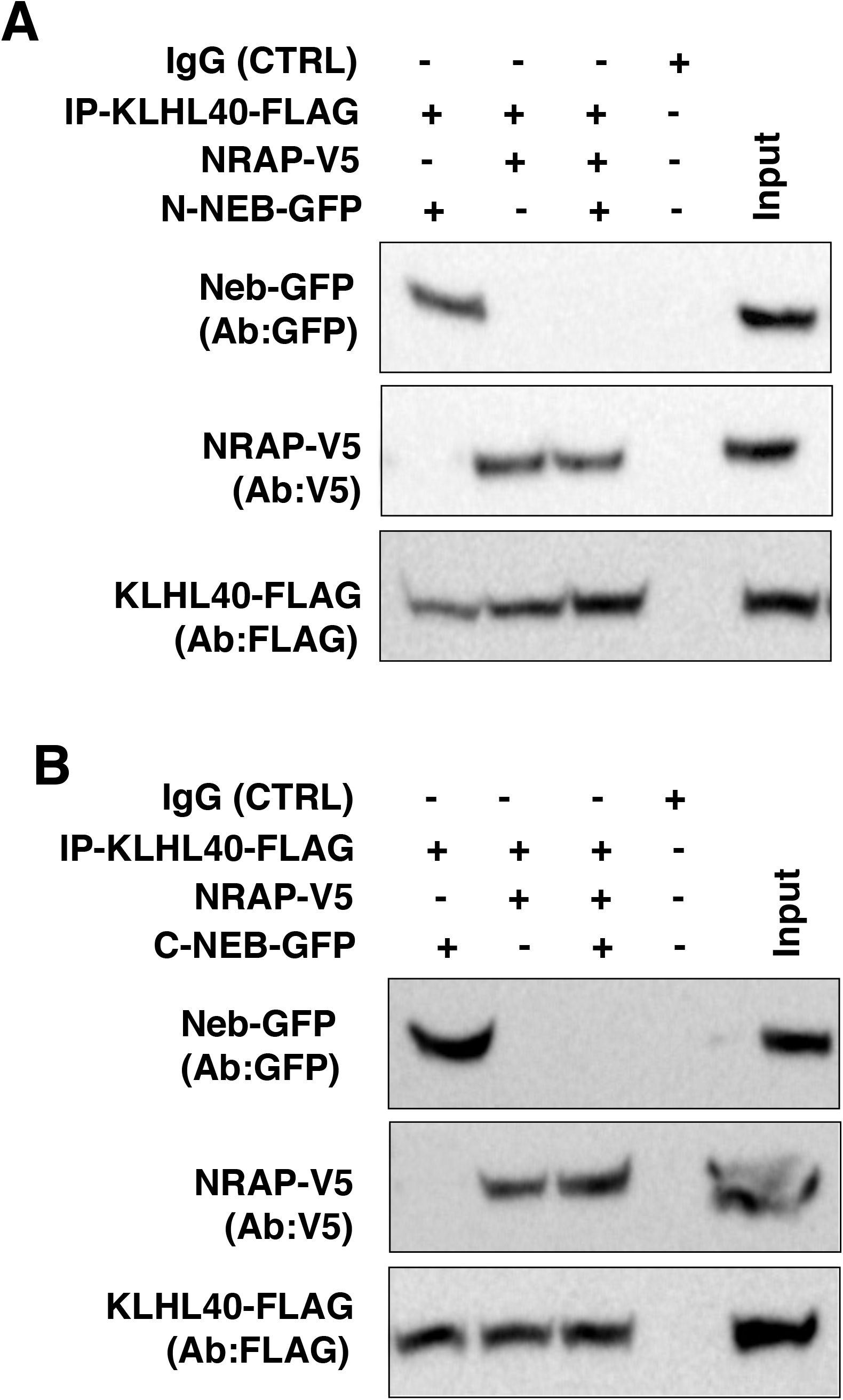
NRAP inhibits KLHL40-NEB interaction. KLHL40 and NRAP or nebulin were overexpressed in C2C12 cells (A) Co-expression of KLHL40 and N-terminal nebulin (N-NEB) or NRAP in C2C12 myoblasts revealed protein-protein interactions between KLHL40/N-NEB or KLHL40-NRAP. In the presence of nebulin, KLHL41 demonstrated preferential binding to NRAP (B) Similarly, overexpression in C2C12 and Western blot analysis revealed that KLHL40 interacts with C-terminal nebulin (C-NEB), and this interaction is blocked in the presence of NRAP.

### Downregulation of NRAP improves structure and function of *Klhl41* deficient zebrafish

Our previous studies have shown *Klhl41* deficient fish recapitulates clinical and pathological hallmark of nemaline myopathy as observed in human patients (15). Therefore, we hypothesized that reducing NRAP levels in KLHL41 deficiency may rescue the pathophysiology observed in *Klhl41*-related nemaline myopathy. To test if reduced NRAP expression KLHL41-NM zebrafish model may rescue the structural and functional deficits in NM, we knocked-down *nrap* in control and *klhl41* morphant fish using antisense morpholino approach. Different concentration of *nrap* morpholino (ex2-in2) (1, 2.5 and 5 ng) was co-injected with *klhl41* morpholinos or control morpholino (Fig. 5A). With increasing concentration of *nrap* morpholino, reduced amounts of *nrap* mRNA were observed in control fish (Fig. S1). No significant changes in skeletal muscle function was observed in control fish at low morpholino concentrations (1.0 and 2.5 ng) (Fig. 5A). At higher morpholino concentration (5.0ng), a mild decrease in swimming behavior was observed in the control fish. Therefore, we used lower concentration of morpholino (1.0 and 2.5 ng) to downregulate *nrap* in control and *klhl41* morphant fish. No improvement in muscle function was observed in *klhl41* morphant fish on injection with lowest morpholino (1.0). Injections with increased amount of morpholino resulted in a significant improvement in muscle function in *klhl41* zebrafish as analyzed by the swimming behavior (Fig. 5B). We previously showed that Klhl41 deficient fish exhibit highly disorganized skeletal muscle with accumulation of nemaline bodies. Electron microscopy evaluation showed while reduction of NRAP (2.5 ng) in control fish did not result in any obvious defects in skeletal muscle ultra-structure, injection of *nrap* MO in *klhl41* morphant fish resulted in an overall improvement in sarcomeric organization as evident by the presence of mature myofibrils (Fig. 5B). Moreover, no nemaline bodies were observed in *klhl41* morphant fish that were co-injected with *nrap* morpholino. These data show that downregulation of *nrap* in KLHL41 deficiency leads to an improvement and structure and function of skeletal muscle.

**Figure 5.**
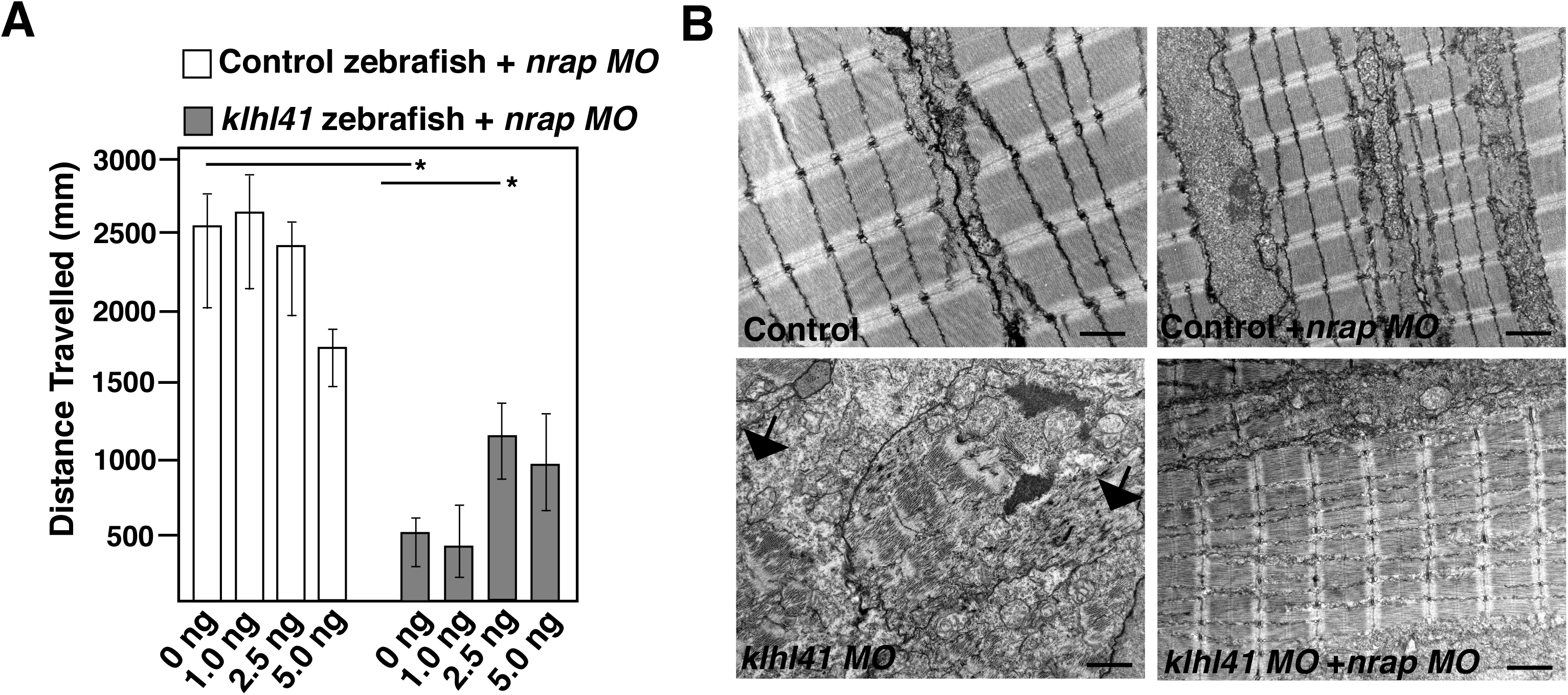
Downregulation of NRAP levels rescues the pathophysiology of KLHL41-NM. Different amount of morpholino targeting zebrafish *nrap* were injected in control or *klhl41* morphant fish (A) Quantification of the swimming behavior in larval fish (3dpf). (B) Skeletal muscle ultrastructure of control and *klhl41* morphant zebrafish targeted with *nrap* morpholino (3 dpf); Scale bar: 500nm.

## DISCUSSION

The UPS pathway is highly conserved in vertebrates and a critical regulator of protein turnover that is required for normal functioning of skeletal muscle. However, little is known regarding the mechanism of protein-turnover dynamics during muscle growth and impact of this process on skeletal muscle development and maintenance. In this work, we provide evidence that KLHL41 regulates sarcomere assembly through ubiquitination and proteasome-mediated degradation of the thin filament chaperone, NRAP. The degradation of NRAP that act in the early steps of sarcomere assembly would comprise a timely and effective negative feedback mechanism to restrain abnormal accumulation of NRAP, thereby regulating the dynamics of myofibril formation. This regulation is critical for disease pathogenesis as abnormal accumulation of NRAP results in myopathic muscle. Therefore, our studies identify NRAP as a potential therapeutic target in NM and uncovers an intriguing cross-talk between ubiquitin-proteasome system and sarcomere assembly process.

We show that KLHL41 interacts with NRAP through C-terminal Kelch domain, and this interaction results in ubiquitination and subsequent degradation of NRAP. Studies have previously identified KLHL41 as an interactor of NRAP, however, functional implications of these interactions in skeletal muscle development and function were not known (51). Our study provides substantial evidence for the destabilizing effect of KLHL41 on NRAP by ubiquitination-mediated proteasomal degradation suggesting selective degradation of proteins may act as a mechanism to regulate the dynamics of sarcomeric proteins that play specific roles during skeletal muscle development. This mode of regulation of skeletal muscle development by ubiquitination is emerging as a critical regulatory mechanism to control protein dynamics and levels. Muscle-specific knockout of the 26S proteasome protein, *Rpt3*, result in significant deficit in muscle growth and force generation in mice (52). Previous studies have shown that UPS-mediated degradation of key myogenic proteins such as Pax7 and MyoD is required for muscle differentiation (53, 54). During muscle differentiation, myoblasts produce increased levels of reactive oxygen species (ROS) leading to an elevated level of oxidized proteins that may negatively affect the muscle differentiation (55). Removal of such proteins by ubiquitin-proteasome system ensures effective muscle differentiation. Interestingly, SMN, mutations, which results in spinal muscular atrophy, is required for normal splicing of ubiquitin-like modifier activating enzyme 1 (UBA1). Low levels of SMN in the disease state results in aberrant splicing of UBA1 and subsequent accumulation of β-catenin. Pharmacological inhibition of β-catenin ameliorated neuromuscular pathology in animal models of SNM deficiency (56). These studies not only demonstrate the extensive regulation of different molecular processes leading to muscle formation by the ubiquitin-proteasome system but also uncover the therapeutic potential of targeting these processes in neuromuscular diseases.

Nemaline Myopathy is a rare skeletal muscle disorder that is caused by mutations in 11 genes, most of which encode for thin filament proteins. We have previously identified mutations in a gene encoding CUL3 E3 ligase adapter, KLHL41, that result in a severe form of nemaline myopathy (15). Mutations in genes encoding closely related Kelch family members, KLHL40 and KBTBD13, also result in recessive and dominant forms of nemaline myopathy, respectively (16, 28). Skeletal muscle biopsies of these patients demonstrate extensive sarcomeric disorganization and abnormal accumulation of protein aggregates or nemaline bodies suggesting a potential defect in the protein turn over process. KLHL41-mediated ubiquitination plays dual roles in regulation of sarcomeric proteins. A previous study has shown that autoubiquitination of KLHL41 is critical for stabilization of nebulin, the major structural constituent of thin filaments in mature myofibers (33). Our work demonstrates that KLHL41 is critical for ubiquitination and subsequent degradation of thin filament chaperone, NRAP, a step that ensures that dynamics of proteins on maturing premyofibrils is maintained.

Our work points to a tightly controlled mechanism by which KLHL41 binds to NRAP and promotes its ubiquitination and subsequent degradation by UPS. Consequently, an increase in NRAP protein is observed in KLHL41 deficiency. The *in vivo* relevance of increased NRAP levels on muscle pathology is evident from *mylz:NRAP-gfp* transgenic zebrafish that exhibit structural and functional deficient in skeletal muscles. While the abnormal accumulation of NRAP has been previously reported in skeletal muscle biopsies in nemaline and myofibrillar myopathy, the functional significance of this abnormal accumulation in disease pathology was not known (57). Our studies suggest that pathologically increased levels of NRAP could be contributing to muscle dysfunction in NM and related forms of myopathies. We further demonstrate NRAP exhibit high affinity for KLHL41 and related family member, KLHL40, and prevents KLHL40-NEB interaction that is required to form mature myofibrils. We speculate that *in vivo*, abnormal accumulation of NRAP may lead to sequestration other muscle proteins that are required for mature myofibril formation and further studies may be able to identify those factors. Previous work has shown that nemaline bodies are formed by different mechanisms and may affect skeletal muscle pathology differently (58). Therefore, identification of different sarcomeric proteins and mechanism contributing to protein aggregates is crucial for a precise understanding of disease biology in NM (Figure 6). Down-regulation of NRAP resulted in a significant improvement in skeletal muscle pathophysiology in *klhl41* deficient zebrafish thus providing direct evidence for reduction of NRAP as a potential therapeutic strategy in KLHL41-related nemaline myopathy. While complete loss of NRAP may lead to skeletal muscle defects, partial downregulation of NRAP may provide an improvement in KLHL41 and related myopathies. Of note, two recent studies have described loss of function mutations in *NRAP* as the cause of underlying disease pathology in dilated cardiomyopathy (59, 60). As asymptomatic sibling also shared the same genotype as the proband, further studies are needed to conclude the pathogenicity of NRAP in human disorders. Downregulation of NRAP mRNA by morpholinos was not able to completely rescue structural and functional deficits observed in KLHL41 deficiency. Future design of therapeutics on targeting abnormal protein rather than mRNA to reduce protein amounts may further provide alternative therapeutic strategy without affecting the basal NRAP transcript levels that are required for normal muscle function. Finally, studies on other *in vivo* targets of KLHL41-Cul3 may provide new insights into suitable pathways and targets that may lead to better therapeutic design in combination therapies based on NRAP downregulation.

**Figure 6.**
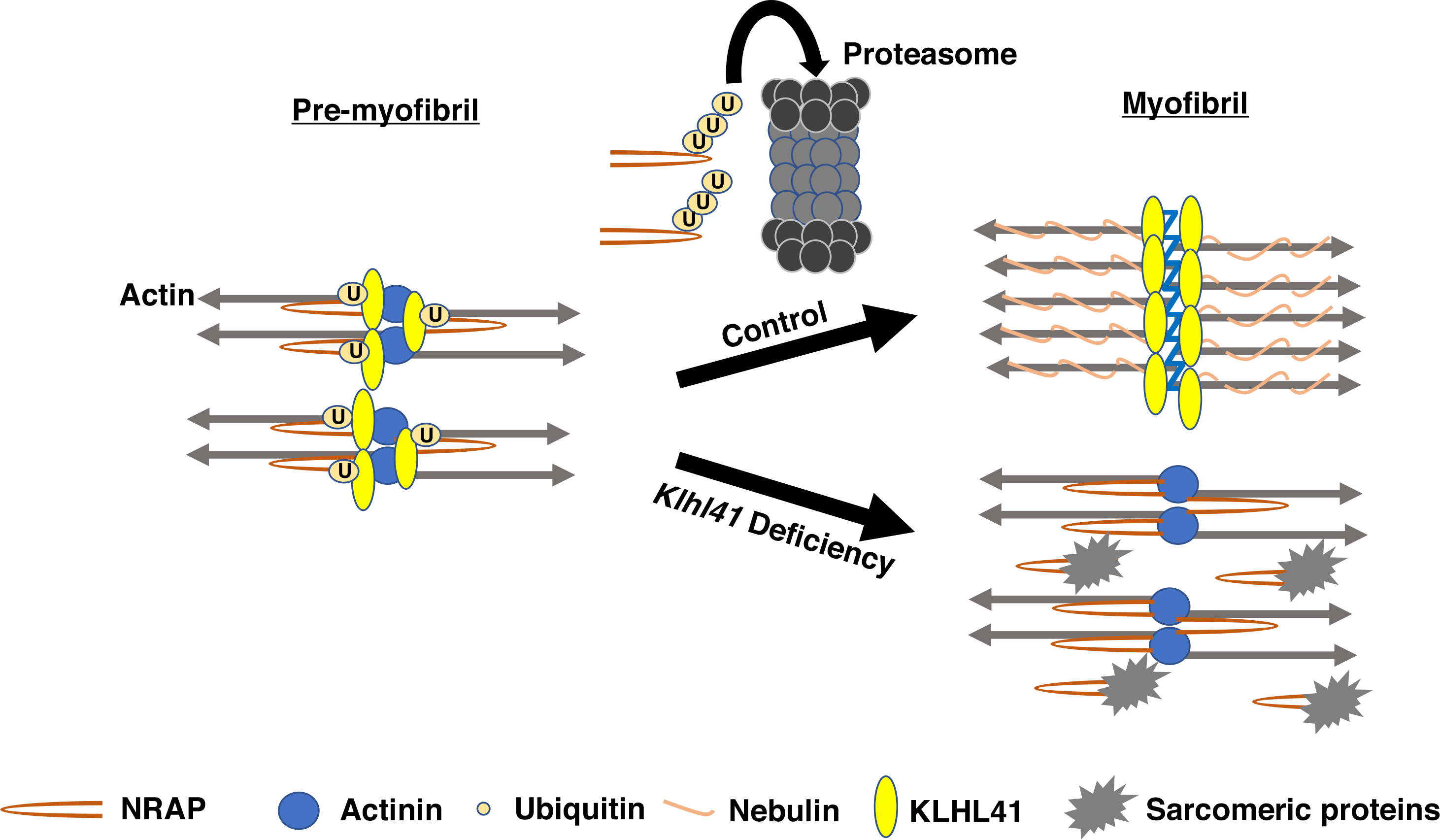
Model of KLHL41 function during sarcomere Assembly. KLHL41 binds to NRAP in pre-myofibrils, and this interaction leads to ubiquitination and subsequent degradation of NRAP by proteasomes. In control skeletal muscle, removal of NRAP from developing pre-myofibrils results in their fusion and formation of mature myofibrils. In the absence of *Klhl41*, abnormal accumulation of NRAP contributes to sequestration of sarcomeric proteins that are otherwise required to form mature myofibers leading to weak muscles.

In summary, KLHL41 acts at the apex position of myofibril maturation by disassembling NRAP by ubiquitination mediated proteolytic degradation with subsequent recruitment and stabilization of nebulin on growing thin filaments thus leading to functional sarcomere assemblies. These studies further propose a paradigm that degradation of skeletal muscle chaperones is required to restrict their function to a narrow temporal window. The identification of a role of KLHL41 in coordinating the premyofibril-myofibril transition offers a unique opportunity to explore the physiological and pathological basis of nemaline myopathy.

## Materials and Methods

### Zebrafish lines

Fish were bred and maintained using standard methods as described (42). All procedures were approved by the Institute Animal Care and Use Committee. Wild-type embryos were obtained from Oregon AB line and staged by hours (h) or days (d) post fertilization at 28.5°C. Zebrafish embryonic (0-2 days post fertilization) and larval stages (3-5 dpf) have been defined as described previously (43).

### Yeast Two-Hybrid

The yeast two-hybrid screen was performed by Hybrigenics S.A.S., Paris, France. Human *KLHL41* cDNA (C-terminal domain) was cloned into pB27 as a C-terminal fusion to LexA (N-LexA-KLHL41-C) and used as a bait for screening a human adult and fetal skeletal muscle cDNA library. The LexA bait construct was used to screen 61 million clones (6-fold coverage of the library). The prey fragments of the positive clones were confirmed for interaction, amplified by PCR, sequenced and identified using the Genbank database (NCBI).

### Time-Resolved FRET Assay (HTRF)

Full-length human KLHL41 proteins was expressed as GST-fusion tag protein in *E.coli*, BL21. Human NEB (aa 5647-5849) and NRAP (aa 1471-1591) were expressed as MBP-HA fusion proteins in *E.coli*, BL21 (Hybrigenics, S.A.S., Paris, France). Cells transfected with fusion plasmids were grown to an optical density of 0.8 (wavelength: 600nM) and induced with IPTG. Cell lysates were prepared by sonication and purified using affinity chromatography. Sonication buffer used for GST tag-fusion protein contained 1X PBS (pH 7.4), 1% Triton X-100, 10% Glycerol. For KLHL41-GST, Glutathione Sepharose 4B (GE Life Sciences USA) was used for affinity purification, and bound fractions were eluted in the presence of 40mM glutathione. The sonication buffer used for MBP tag-fusion protein contained 20mM PBS (pH 7.4), 200mM NaCl, 0.5% Triton X-100, 1mM EDTA, 2mM DTT and 10% Glycerol. Amylose affinity resin (New England Biolabs, USA) was used for affinity purification, and bound fractions were eluted in the presence of 10mM maltose. For HTRF assay, dilutions of both interactor proteins and antibodies were freshly prepared and incubated together for 2hr at 4°C. Following this 10μL anti-GST-EuK and/or anti MBP-d2, antibodies were added (1:400 dilution). The interaction between two proteins was detected by fluorescence transfer (excitation at 337 nm, emission at 665 nm). The emission at 620 nm occurs regardless of the interaction and used for normalizing the assay. The interaction between two proteins was quantified as DeltaF (%) as:

> DeltaF (%) = 100 X (ratio_sample_ - ratio_background_)/ratio_background_

where “ratio” is the 665/620 fluorescence ratio, “sample” is the signal in the presence of protein interactors and antibodies, whereas, “background” denotes the HTRF antibodies only. Fusion tags without interactor proteins were used as negative controls.

### Co-immunoprecipitation

Human KLHL41-pEZYFLAG, human NRAP-V5, C-terminal NEB-GFP and N-terminal NEB-GFP were used for co-immunoprecipitation assay in C2C12 cells. Equimolar concentration of KLHL41-pEZYFLAG, NRAP-V5 or NEB-GFP and control empty plasmids were transfected in C2C12 cells. 48 hours post transfection, cell lysates were prepared in RIPA buffer and precleared with mouse IgG and Protein A/G agarose beads for 1 hr at 4°C. V5-agarose (Thermo Fisher Scientific) or FLAG-agarose (Sigma, St. Louis, MO, USA) beads were incubated with the supernatant (4°C, overnight) and washed three times with PBS and once with high stringency buffer (300mM NaCl + PBS buffer). For all experiments, two negative controls consisted of a sample lacking the primary antibody (beads) and a sample incubated with IgG. Resulting protein complexes were eluted in 1X LDS buffer (Thermo Fisher Scientific) and analyzed by SDS-PAGE and Western blotting using KLHL41 antibody, (1:500, AV38732, Millipore Sigma, USA), FLAG antibody (1:500, F1804, Millipore Sigma), V5 antibody (1:500, R960CUS, Thermo Fisher Scientific, USA) and GFP antibody (1:250, sc-9996, Santa Cruz Biotechnology, USA).

### Whole Mount phalloidin staining

Zebrafish larvae (3 days post fertilization) were fixed in 4% PFA overnight at 4°C, then washed as follows: 2 × 10 min in PBS, 2 × 10 min in PBS-T (0.1% Tween-20), 1 × 60 min in PBS-TR (2% Triton X), and 2 × 5 min in PBS-T. Embryos were blocked in PBS-T containing 5% goat serum for 1 hour at RT and incubated with phalloidin (1:40, Thermo Fisher Scientific, A12379). Subsequently, larvae were washed for 4 × 15 min in PBS-T, mounted in 70% glycerol and visualized using a Perkin Elmer UltraVIEW VoX spinning disk confocal microscope.

### C2C12 myoblasts culture

Mouse C2C12 cells were cultured in a growth medium consisting of DMEM supplemented with 20% fetal bovine serum. To create *Klhl41* knockout lines, Cas9-GFP plasmid was co-transfected with sgRNAs targeting exon1 of mouse *Klhl41* gene. 24 hours post transfection, GFP positive cells were selected by FACS, and single cells were plated in each well of a 96 well plate in the C2C12 conditioned media. Clonal expansion was performed, and loss of function mutations were validated by Sanger sequencing and Western blot analysis.

### Ubiquitination studies

C2C12 cells were transfected with different amounts of KLHL41-pEZYFLAG, NRAP-V5 and HA-ubiquitin plasmids (44). MG132 (10uM) was added at 40 hours post transfection, and cells were harvested at 48 hours post transfection. Cell lysates were prepared in RIPA buffer. To reduce the non-specific binding during immunoprecipitation, supernatant was incubated with mouse IgG and Protein A/G agarose beads for 1 hr at 4°C followed by overnight incubation with V5-agarose at 4°C. Beads were washed with 1XPBS buffer (3 times) and boiled in 1X LDS sample buffer to elute proteins. SDS-PAGE and western blotting were performed as previously described (45). To analyze proteasome mediated degradation of NRAP, control or *Klhl41* knockout C2C12 myoblasts were grown in the presence or absence of proteasome inhibitor (MG132, 10μM) for 48 hours, and proteins were analyzed by Western blot analysis. For cycloheximide chase assay, control or *Klhl41* knockout C2C12 cells (5 × 10^5^) were plated in 35 mm plates. After 12 hr, replace with fresh medium containing 100μg/ml cycloheximide. Samples were collected at different time points; protein lysates were prepared, and Western blotting was performed.

### NRAP overexpression and knockdown in zebrafish

Human *NRAP* was integrated in to zebrafish genome utilizing the Tol2 transposon-based system (46). A GFP reporter-protein was linked to the coding sequence of *NRAP* under the control of *mylz*, a skeletal muscle specific promoter (*mylz:NRAP-gfp*) using Gateway multisite cloning. 25 ng of the transgenic plasmid was co-injected with 10 ng of transposase mRNA in to 1-cell zebrafish embryos. F^0^ zebrafish were raised and crossed with wild-type zebrafish (AB strain). Germline transmission of the *NRAP* transgene in F^1^ generation was confirmed by PCR-based genotyping using primers for GFP: forward, 5‘-AAGCTGACCCTGAAGTTCATCTGC-3’, and reverse, 5‘-CTTGTAGTTGCCGTCGTCCTTGAA-3’. Additionally, the expression of hu-NRAP-GFP protein was confirmed by immunoblotting with a GFP antibody at 3 dpf that showed ~3.4 folds upregulation of the transgene compared to wild-type controls

Two antisense MO, targeting the exon-intron junction of zebrafish *nrap* gene (NM_001324547) were designed to knockdown the zebrafish *nrap* transcript. The MO sequences were designed to target exon-intron junctions; exon2-intron2: 5‘-TGTGCCAGTTCTGAAAGAGAAAAGA-3’ and exon6-intron6: AACATAAATCCTTACATCACTGGCC.

MO against human *β-globin*, which is not homologous to any sequence in the zebrafish genome by BLAST search, was used as a negative control for all injections (5‘-CCTCTTACCTCAGTTACAATTTATA-3’). *klhl41* morpholino sequences have been described previously (15). Sequences of PCR primers used for zebrafish *nrap* gene is: Forward, 5‘-CGCCTTGCTGTTCCTATACC-3’ and Reverse, 5‘GCCTCTGATTGCTTCTTTGC3‘. MOs were dissolved in 1X Danieau buffer with 0.1% phenol red and 1–2 nl (1–10 ng) injected into the yolk of 1-cell stage embryos.

### Zebrafish locomotion assay

Zebrafish swimming behavior was quantified by an infra-red tracking activity monitoring system (DanioVision, Noldus, Leesburg, VA, USA). Control, transgenic or morphant larval zebrafish were placed individually into each well of a 24 well plate in dark for 10 minutes. The activity of these larvae was recorded during a follow-up light exposure of 20 minutes. Four independent blind trials were performed, and mean velocity, total distance and cumulative duration of movement were recorded. Reported values reflect an average of 30-35 larval fish.

### Quantification and statistical analysis

All samples were blinded till final analyses. Data were statistically analyzed by parametric Student t-test (two tailed) and were considered significant when P<0.05. All data analyses were performed using XLSTAT software.

## Acknowledgement

This work was supported by Miles Shore Fellowship for Scholars in Medicine and Brigham and Women’s Hospital Career Development Award and A Foundation Building Strength Grant to VAG.

## Supplementary Data

**Figure S1.**
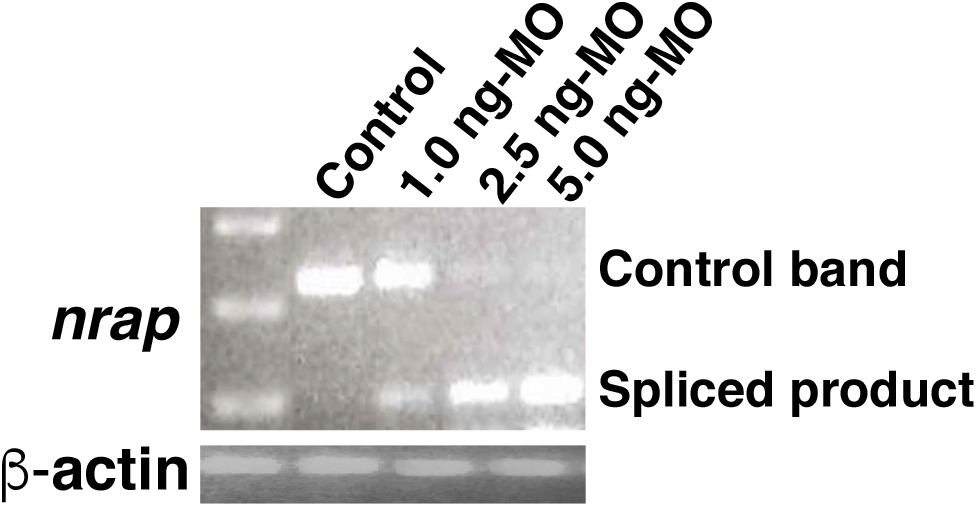
RT-PCR analysis of *nrap*. Control 1-cell zebrafish embryos were injected with different amounts of *nrap* morpholino (ex2-in2). RT-PCR analysis was performed using primers targeting exon1 and exon3 of zebrafish *nrap* gene.

**Table S1.** KLHL41 Interactors in skeletal muscle identified by Yeast-2 Hybrid Screening.

| Interactor (Gene Name) | Function |
| --- | --- |
| Nebulin (NEB) | Major structural constituent of thin filament |
| Nebulin Related Anchoring Protein (NRAP) | Thin Filament Chaperone |
| Proteasome Maturation Protein (POMP) | Essential for 20S Proteasomal Formation |
| Interferon Related Developmental Regulator (IFRD1) | Positive regulator of skeletal muscle differentiation by increasing muscle stem cells |
| RNA Binding Motif Protein 4 (RBM4) | Splicing regulator, overexpression promotes muscle cell differentiation |

